# Revealing intraosseous blood flow in the human tibia with ultrasound

**DOI:** 10.1101/2021.04.12.439472

**Authors:** Sebastien Salles, Jami Shepherd, Hendrik J. Vos, Guillaume Renaud

## Abstract

Intraosseous blood circulation is thought to have a critical role in bone growth and remodeling, fracture healing, and bone disorders. However, it is rarely considered in clinical practice due to the absence of a suitable non-invasive in vivo measurement technique. In this work, we assessed blood perfusion in tibial cortical bone simultaneously with blood flow in the superficial femoral artery with ultrasound imaging in 5 healthy volunteers. After suppression of stationary signal with Singular-Value-Decomposition, pulsatile blood flow in cortical bone tissue is revealed, following the heart rate measured in the femoral artery. Using a method combining transverse oscillations and phase-based motion estimation, two-dimensional vector flow was obtained in the cortex of the tibia. After spatial averaging over the cortex, the peak blood velocity along the long axis of the tibia was measured four times larger than the peak blood velocity across the bone cortex. This suggests that blood flow in central (Haversian) canals is larger than in perforating (Volkmann’s) canals, as expected from the intracortical vascular organization in humans. The peak blood velocity indicates a flow from the endosteum to the periosteum and from the heart to the foot for all subjects. Because aging and the development of bone disorders are thought to modify the direction and velocity of intra-cortical blood flow, their quantification is crucial. This work reports for the first time an in vivo quantification of the direction and velocity of blood flow in human cortical bone.

## INTRODUCTION

Intraosseous blood circulation is thought to play a key role in bone growth and remodeling, bone fracture healing or osteointegration of bone scaffold, and the development of bone disorders and metastasis (1–8). In particular, inadequate bone vascularity (hypo-vascularization or hyper-vascularization) is associated with bone disorders like osteoporosis and osteonecrosis.

In the diaphysis of long bones, osteons are nearly parallel to the axis of the bone. The canal provides a passage for blood vessels and nerve fibers through the hard bone matrix. Most canals contain a single vessel of capillary structure, although wide canals can contain both an arteriole and a venule (7). In addition, the presence of perforating Volkmann’s canals (Figure 1.1) ensures communication between the Haversian canals and circulation between the outer and inner spaces (periosteum and medullary cavity). Human cortical bone contains more Haversian canals than Volkmann’s canals (9). In the cortex of the diaphysis of an adult human long bone, the Haversian canals are nearly aligned with the axis of the long bone (10). Therefore blood in human diaphyseal cortical bone is expected to circulate mainly in the direction of the long bone axis.

The median diameter of Haversian canals in adult human cortical bone was reported between 40 and 100 micrometers, and the density of Haversian canals is close to 10 pores per square millimeter (11–13). A recent study observed blood vessels in human femoral cortical bone with a diameter of 50 +/− 10 micrometers (14). Older investigations reported intracortical vessels with a diameter of 15-30 micrometers (15).

Evidence of the importance of intraosseous blood circulation in bone disorders has been obtained in animal models, most often with invasive techniques (16).

A microsphere technique is generally accepted as the “gold standard” for measuring blood perfusion in bone, in animals (17). It is invasive, as it requires sampling of tissue and embolization of capillaries, and is therefore inapplicable in humans. However, evidence in humans is very sparse because of the absence of a suitable non-invasive technique.

More recently, a transcranial imaging approach based on hybrid optoacoustic and ultrasound bio-microscopy revealed the vascular morphology in murine skulls, although without quantification of blood flow (18). However, this technique is likely inapplicable for non-invasive imaging of human bones due to a very small imaging depth.

In humans, the feasibility of assessing intraosseous blood circulation has been studied with several techniques. Dynamic contrast-enhanced magnetic resonance imaging (MRI) and diffusion-weighted MRI have been proposed to assess marrow perfusion in trabecular bone and were limited to rather large regions of interest (vertebra, femoral head)(19–21). Recently Wan et al. (22) have shown that dynamic contrast-enhanced MRI can assess cortical bone perfusion. PET uses ionizing radiation, and its low spatial resolution (about 5 mm) does not allow the clear distinction between blood flow in cortical bone, marrow, and soft tissues surrounding bone. Even if dynamic contrast-enhanced MRI and PET can provide an absolute estimation of blood flow, the acquisition time is long (up to 1 hour); the outcome relies on a compartmental model and the measurement of an arterial input function. Researchers often choose to measure more straightforward parameters that do not provide an absolute measure of blood flow (20,23). Near-infrared optical methods have also been applied to the assessment of cortical bone perfusion, but they are limited to an investigation depth of about 1 cm and, like PET, have low spatial specificity (24). It is worth noticing that none of the existing techniques to measure intraosseous blood circulation can determine the direction of blood flow. Moreover, while these techniques have been used for research, they are not used for routine clinical diagnosis of cortical bone perfusion.

Ultrasound imaging is relatively inexpensive, does not produce ionizing radiation, offers good spatial resolution and excellent temporal resolution, and has the potential to investigate deeper tissues compared to optical techniques. Ultrasonography provides a direct and absolute measurement of the velocity and direction of blood flow (25,26). Nonetheless, clinical ultrasound scanners currently rely on a major assumption during image reconstruction: the speed of sound in the region of interest is assumed to be uniform. This assumption is valid for soft tissues, but it does not hold for bone. As a result, conventional ultrasonography fails to evaluate blood flow in cortical bone and marrow; only the vascularization of the periosteum (the membrane that covers the outer surface of bones, as shown in Figure 1) could be assessed (27,28).

**Figure 1:**
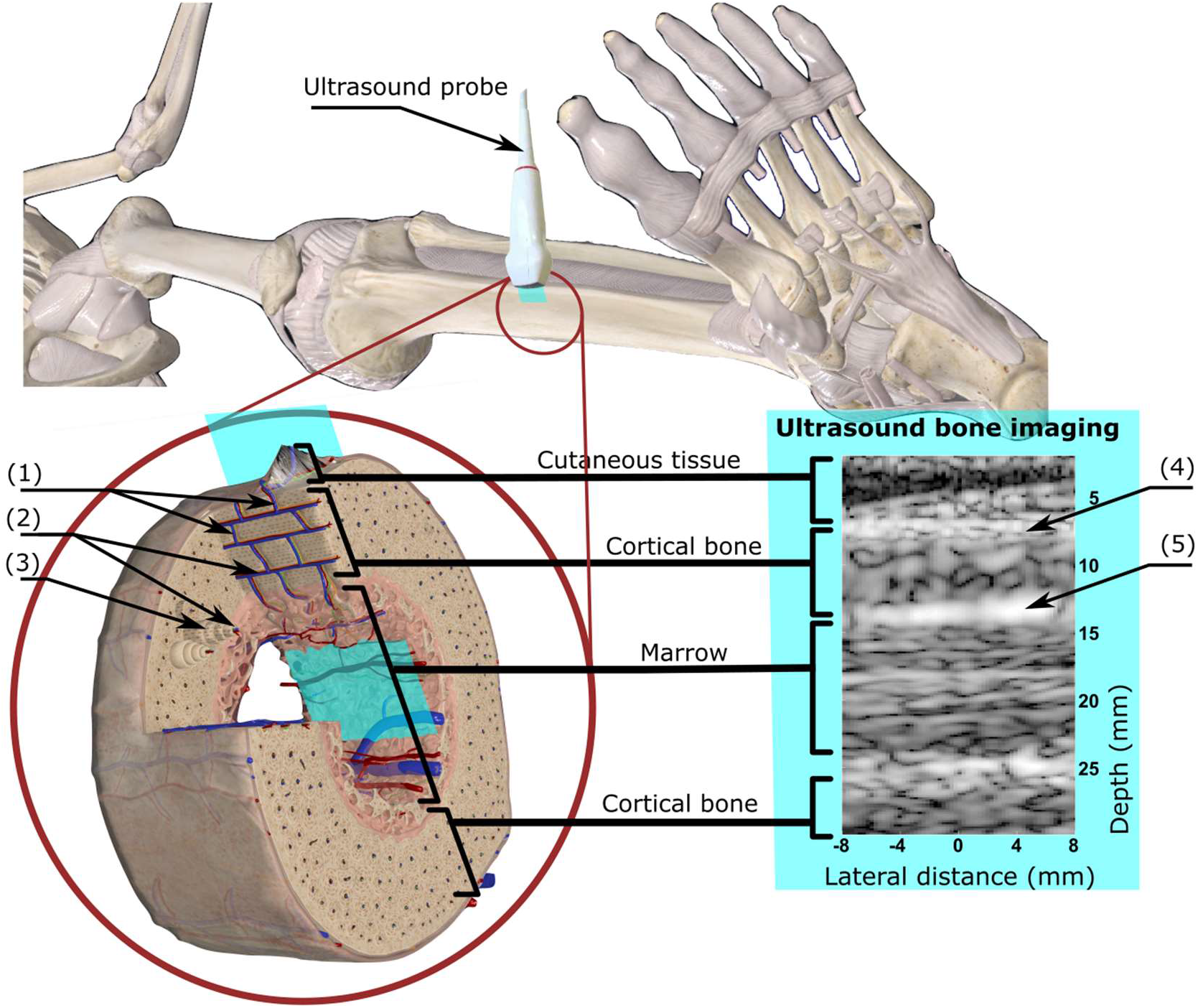
Ultrasound imaging of the bone cortex at the tibia. This figure shows a three-dimensional schematic representation of bones in the leg, the probe positioning on the tibia, the image plane, and the corresponding ultrasound image. The ultrasound probe was placed on the medial surface of the tibia, in the middle of the diaphysis. The blue plane indicates the ultrasound image plane. Three distinct layers are visible in the ultrasound image, namely cutaneous tissue, cortical bone, and the marrow. Haversian canals (2) provide a passage for blood vessels and nerve fibers through the hard bone matrix. In addition, the perforating Volkmann’s canals (1) ensure communication between Haversian canals. Blood can therefore circulate between the periosteum and the endosteum. Note that the three-dimensional schematic representation of the vascular organization in the diaphysis of a long bone depicts a cortex with a small thickness (less than 1 mm, while the tibial cortical thickness is about 5 mm). Modified excerpt from Complete Anatomy ‘20 with permission from 3D4Medical (www.3d4medical.com). (1): Volkmann’s canal, (2): Harversian canal, (3): Osteon, (4): Periosteum, (5): Endosteum

We present an approach that provides a directional and quantitative measurement of blood circulation in human cortical bone for the first time, in vivo, using ultrasound imaging.

## MATERIALS AND METHOD

### Refraction-corrected ultrasound imaging of bone

In this work, we used synthetically-focused ultrasound imaging consisting of a coherent summation of images with low contrast resolution, obtained from different insonification angles, into one image with high contrast resolution. Low-resolution images can be achieved in different ways, typically by transmitting either plane or spherical waves(29,30). As shown in the Supplemental figure 1, we used the transmission of multiple steered plane waves. Unlike conventional focused-beam scanning, this technique allows synchronous measurement of blood flow in the entire region of interest(31).

In this study, an ultrasound probe was placed on the medial surface of the tibia of volunteers to generate a longitudinal 2D image of the diaphysis. Relying on previous work (32), refraction-corrected image reconstruction was accomplished. Unlike conventional medical ultrasonography, which assumes a homogeneous medium during image reconstruction, our approach describes the scanned region as a layered medium. Three layers are considered, namely cutaneous tissue, cortical bone, and marrow.

The compressional wave-speed in cortical bone is more than double the wave-speed in soft tissues. The resulting change in propagation direction (refraction) as ultrasound waves traverse the outer or inner surface of the bone cortex must be taken into account for accurate image reconstruction and blood velocity estimation. In addition, cortical bone exhibits wave-speed anisotropy that was also accounted for in our reconstruction method using a 3-parameter weak anisotropy model (33,32). The wave-speed values previously measured in healthy volunteers were used for all healthy volunteers in this new study. More details are provided in the Methods section. Then, after accurate calculation of round-trip travel times of ultrasound waves, a delay-and-sum approach is applied for reconstructing the images.

### Blood flow estimation in cortical bone

In order to capture blood flow, the procedure to generate one image of the cortex will be repeated 100 times per second, during a continuous examination of 4 seconds. With the ultrasound technique used in this study, we achieved a spatial resolution close to 1.5 mm in the image of the bone cortex. Consequently, the pores and blood vessels in the cortical bone were not resolved in the ultrasound image. Instead, an ultrasound image of the bone cortex shows speckle, as seen in Figure 1.

During the 4 seconds of an acquisition, the movement of erythrocytes through blood vessels causes temporal fluctuations in the image. However, relative motion between the hand-held ultrasound probe and the tibia causes small and slow fluctuations of image intensity as well. These blood-unrelated fluctuations must be removed to allow quantification of blood velocity. In general, blood-unrelated fluctuations have a velocity smaller than fluctuations caused by blood flow (Supplemental figure 1c-d.) and high spatial correlation.

Singular value decomposition (SVD) was used to extract the time-variant component in the ultrasound image caused by blood flow (34)(Supplemental figure 1b). Next, blood-related fluctuations in the image were analyzed to calculate three flow metrics: a flow metrics proportional to the volume of blood in motion (“power Doppler”) (35) and the two components of blood velocity in the 2D image, i.e. the axial velocity (blood circulating in the direction of the long bone axis) and the radial velocity (trans-cortex blood circulation in a direction perpendicular to the long bone axis). Power Doppler has the advantage of being robust to noise, but it is relatively angle independent (35) and provides no absolute measurement of blood flow velocity. Despite their higher sensitivity to noise, several methods are known in the literature for estimating the 2D vector flow mapping (25,26,36). Here, we chose a method combining a phase-based motion estimation and the transverse oscillation technique (37–39) to calculate the axial and radial components of blood velocity at all pixels in the ultrasound image (see Appendix section for more details).

The 4s acquisition procedure was repeated 3 times with repositioning of the probes on 5 healthy volunteers. A continuous recording of 4 seconds ensured that at least 3 cardiac cycles were observed each time. Each subject was laid down on a hospital bed.

### Blood estimation in the superficial femoral artery

To compare the pulsatility of the blood flow measured in tibial cortical bone with that in the main arterial input in the lower leg, the ultrasound examination was performed simultaneously at the tibia and at the superficial femoral artery, using two hand-held ultrasound probes operated by one research ultrasound system. Flow quantification in the superficial femoral artery was performed with conventional Doppler ultrasound imaging (see Methods section for details).

Peak flow velocity and heart rate were assessed in the femoral artery (Figure 2) for comparison with the periodicity of blood flow estimated in tibial cortical bone. Because the power Doppler signal is more robust to noise, we used this hemodynamics parameter to estimate the heart rate.

**Figure 2:**
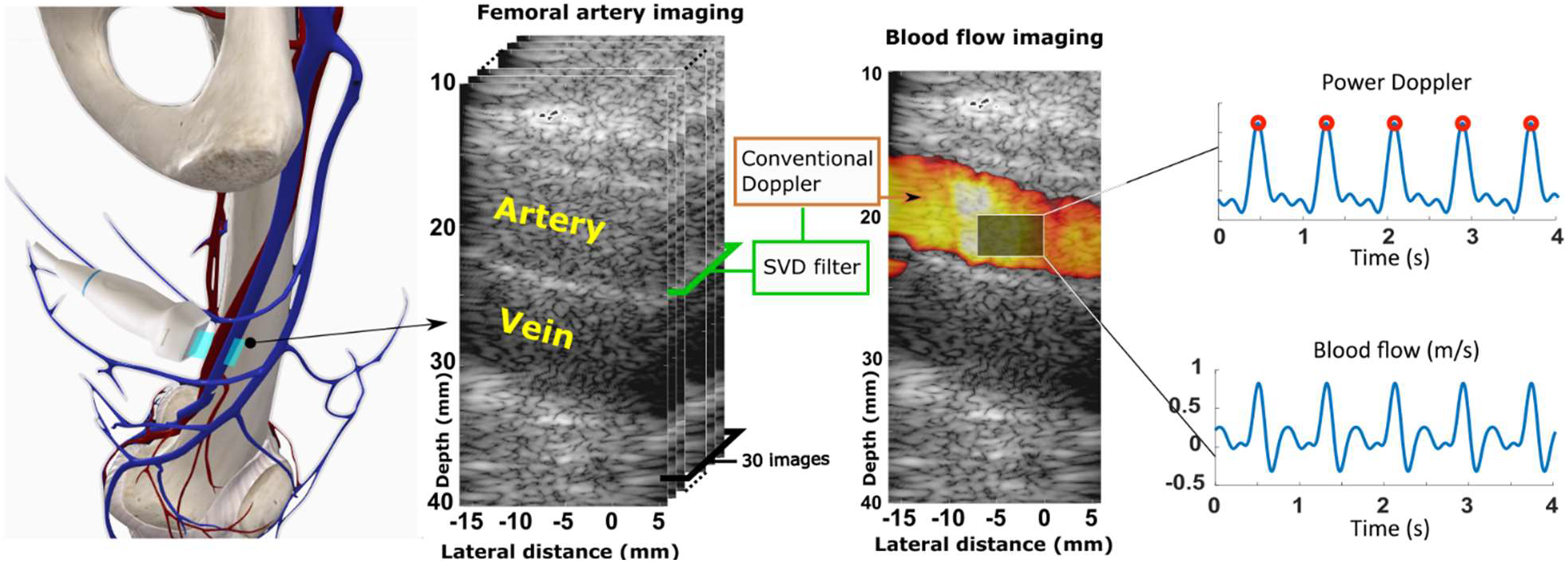
Blood flow imaging in the superficial femoral artery. 400 ultrasound Doppler quantifications were acquired at a rate of 100Hz. Each quantification was obtained with 30 identical titled (20°) plane-wave transmissions at a pulse repetition rate of 5000 Hz. After SVD filtering, a conventional Doppler technique was applied to each ensemble of 30 images. The two blue curves show the power Doppler signal and blood flow velocity along the artery’s axis. The red circles indicate the power Doppler signal peaks and were used to measure the heart rate. Modified excerpt from Complete Anatomy ‘20 with permission from 3D4Medical (www.3d4medical.com).

#### Human subjects

Five healthy subjects (Table 1) were recruited following approval from the local medical ethics committee of Erasmus MC University Medical Centre Rotterdam, The Netherlands (MEC-2014-611). All participants provided written informed consent.

**Table 1:** Subject age, total number of recorded cardiac cycles and parameters for SVD clutter filtering and transverse oscillations

| Subject | Age | Total number of cardiac cycles | Method parameters |  |
| --- | --- | --- | --- | --- |
| | | | SVD-filter threshold | Transverse oscillations $\lambda_{0x}$ (mm) |
| 1 | 32 | 13 | 50 | 3.5 |
| 2 | 46 | 9 | 40 | 3.5 |
| 3 | 65 | 9 | 50 | 3.5 |
| 4 | 60 | 8 | 40 | 3.5 |
| 5 | 28 | 10 | 30 | 3.5 |

## Results

### Hemodynamics measured in tibial cortical bone

The proposed approach aimed to characterize blood flow in small vessels both in Haversian canals (along the bone axis) and in Volkmann’s canals (perpendicular to the bone axis). Therefore, our analysis provides directional information, which is of interest for better understanding of bone physiopathology.

The measured bone displacement with respect to the probe was inferior to 0.5 mm, with a peak velocity close to mm/s, for all subjects and showed no pulsatility (Supplemental figure 1d). After SVD filtering of these slow fluctuations caused by relative probe-bone motion, the results revealed pulsatile blood flow in cortical bone tissue. Supplemental figure 1c shows that reproducible pulsatility was observed in all hemodynamic parameters at a given image pixel, namely power Doppler, axial velocity, and radial velocity.

The ultrasound image did not resolve the small blood vessels in cortical bone, and cortical bone possesses a rather organized vascularization with two principal flow directions (Haversian and Volkmann’s canals). Because of these two facts, we proposed to spatially average the blood-velocity vector field over the investigated region of cortex (Figure 3), which corresponds to a volume of 5mm x 15mm x 10mm (out-of-plane image thickness). Interestingly, after spatial averaging over the cortex, pulsatile perfusion was observed in tibial cortical bone in all 5 subjects (Figure 3 and Figure 4). Table 2 and Figure 5 summarize the quantitative flow metrics measured for the 5 subjects. The peak blood velocity in the direction of the tibia axis (axial velocity) was 4 to 5 times larger than that across the cortex (radial velocity). For all 5 subjects, the maximum measured blood motion corresponded to blood circulating from the medullary cavity to the periosteum (i.e. centrifugal flow, from the marrow to the outside of the tibia), and from the heart to the foot.

**Figure 3:**
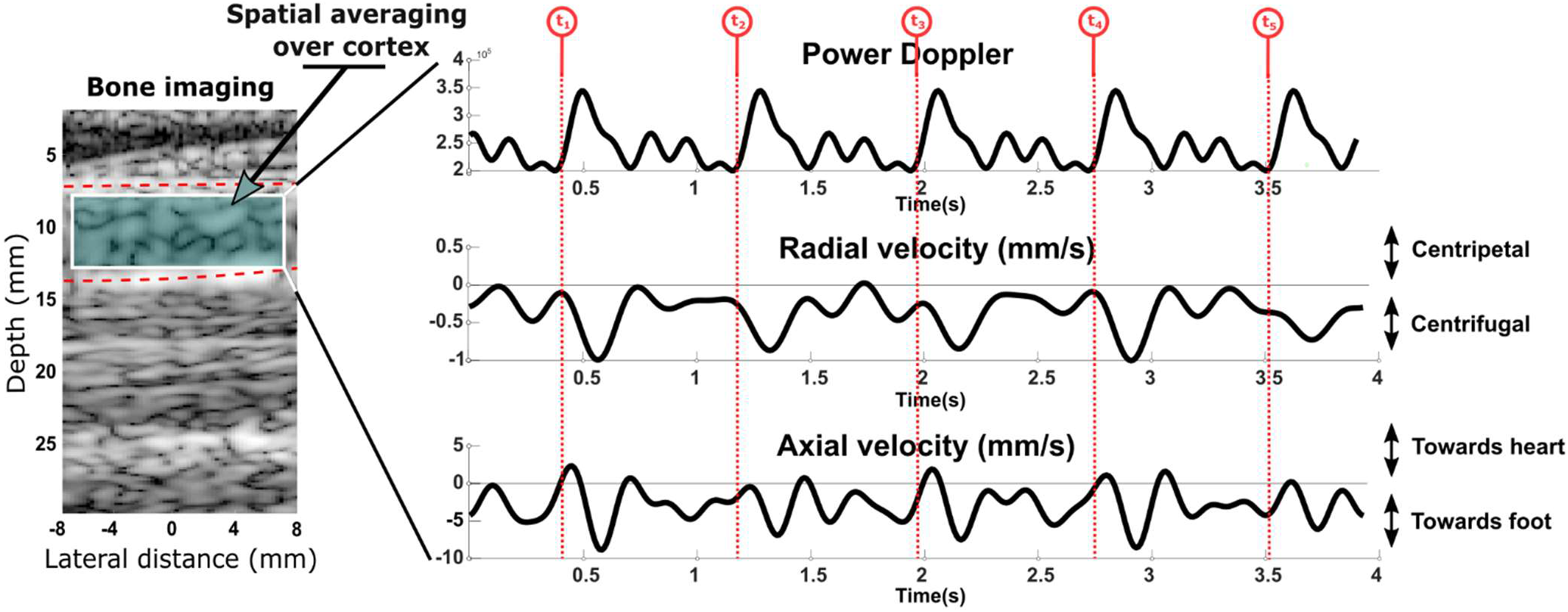
Blood flow in tibial cortical bone measured in subject 1, after spatial averaging over the bone cortex. Pulsatile blood flow in cortical bone tissue is visible in each hemodynamic parameter, namely the power Doppler and the axial and radial velocities. Similar velocity waveforms were observed at the peak flow within 5 consecutive cardiac cycles (t1, t2, t3, t4, t5).

**Figure 4:**
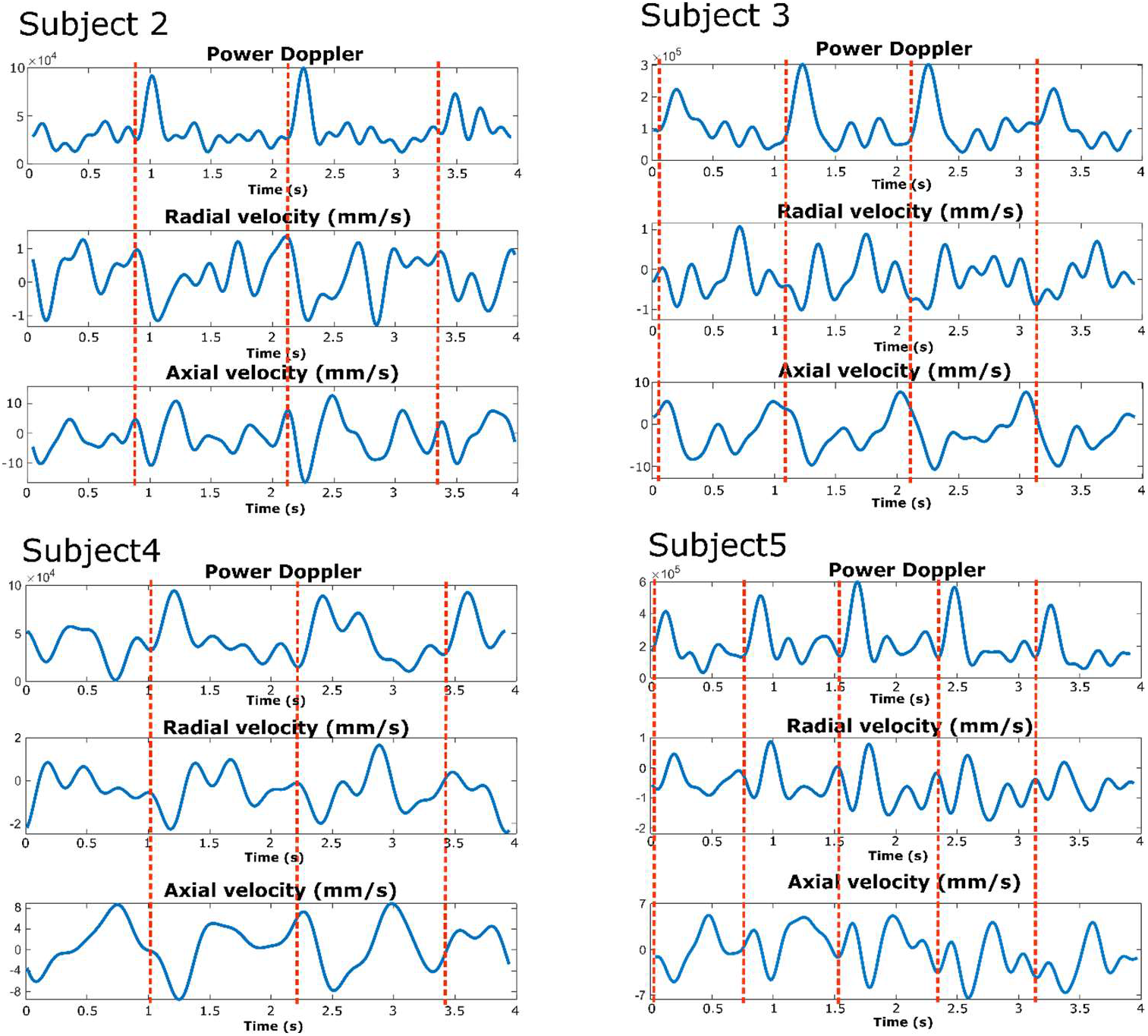
Examples of blood flow measurements in tibial cortical bone obtained in subjects 2-5, after spatial averaging over the bone cortex. Pulsatile blood flow in cortical bone tissue is visible in each hemodynamic parameter, namely the power Doppler and the axial and radial velocities. The red dash line indicated the cardiac cycles.

**Table 2:** The peak negative, peak positive, and temporally averaged values of the axial, and radial blood velocity in tibial cortical bone averaged over the 5 subjects. The sign convention is depicted in Figure 3.

| Blood velocity (mm/s) | Average over 5 subjects | Standard deviation |
| --- | --- | --- |
| Axial negative peak (Long bone axis, towards foot) | -9.9 | 2.3 |
| Axial positive peak (Long bone axis, towards the heart) | 7.9 | 1.6 |
| Axial time average (Long bone axis) | -0.4 | 0.3 |
| Radial negative peak (Centrifugal trans-cortex flow) | -2.3 | 0.7 |
| Radial positive peak (Centripetal trans-cortex flow) | 1.6 | 0.3 |
| Radial time average (trans-cortex flow) | -0.1 | 0.1 |

**Figure 5:**
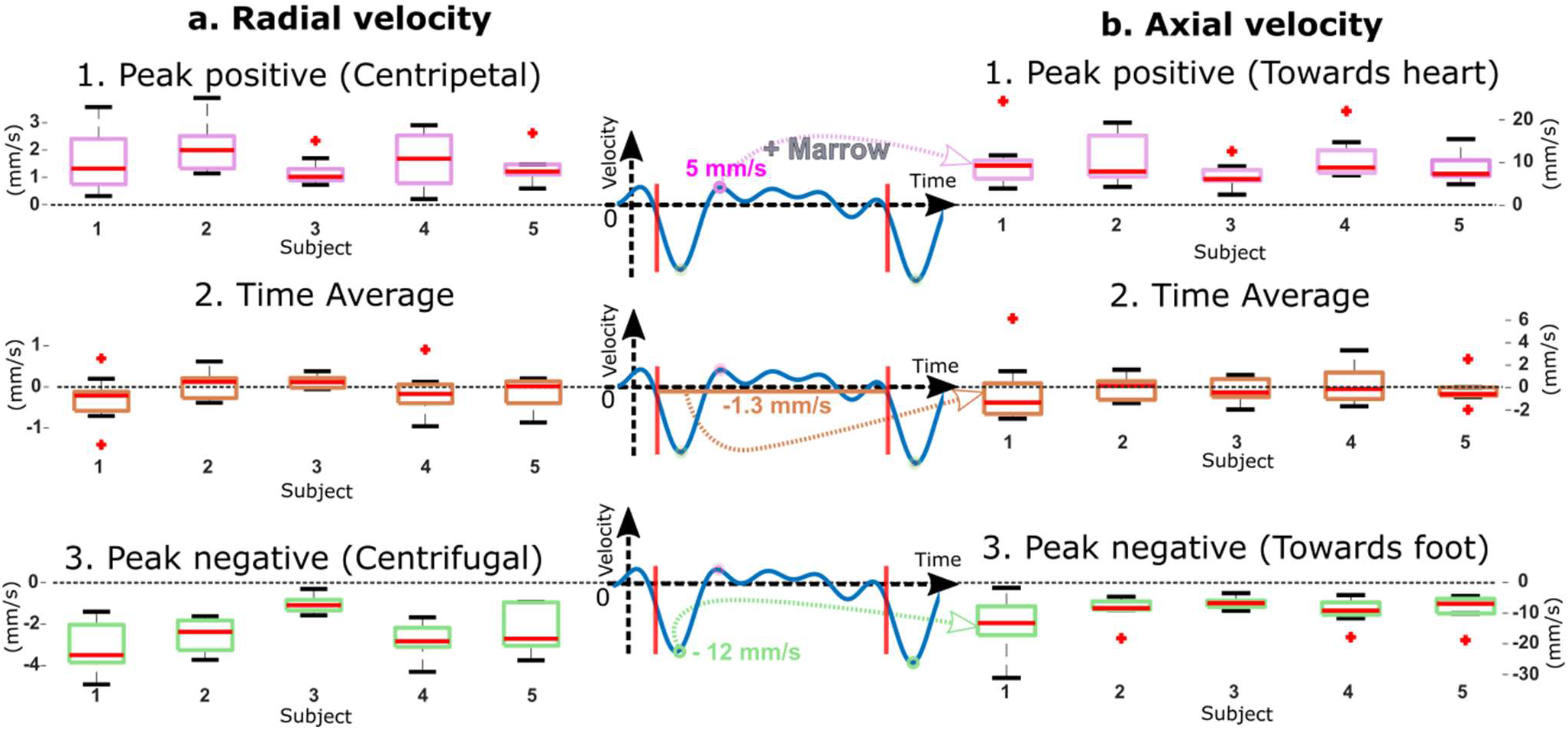
Blood velocity measured in tibial cortical bone for the 5 subjects. The left column (a) shows the radial blood velocity corresponding to blood flow in Volkmann’s canals. The right column (b) shows the axial velocity corresponding to flow circulating in the Haversian canals. Both columns include the peak positive, peak negative, and time-averaged values estimated in multiple cardiac cycles for each volunteer. The middle column shows an example of the axial velocity. The axial (radial) peak positive velocity was found between 5.7 (1.3) mm/s and 9.4 (2) mm/s, and the peak negative velocity between −13.9 (−3) mm/s and −7.7 (−1) mm/s, respectively. In each box, the central mark indicates the median, red crosses indicate outlier values, and the bottom and top edges of the box indicate the 25th and 75th percentiles, respectively. The three identical signals in the middle column are used as an illustration to define the quantification parameters. The axial and radial velocities’ parameter values are also shown in Table 3 and 4, respectively.

The time-averaged radial and axial blood velocities measured in cortical bone (Table 2 and Figure 5) are small (less than 1 mm/s) and show large variabilities. Therefore, no physiological interpretation of these flow metrics can be made. The minimum peak blood velocities were observed in volunteer 3 (Figure 5, Table 3 and 4); the axial and radial velocities reached 8 mm/s and 1 mm/s, respectively (averaged over 12 cardiac cycles). The maximum peak blood velocities were observed in volunteer 1 (Figure 5 Table 3 and 4); the axial and radial velocities reached 14 mm/s and 3 mm/s, respectively (averaged over 9 cardiac cycles).

**Table 3:**
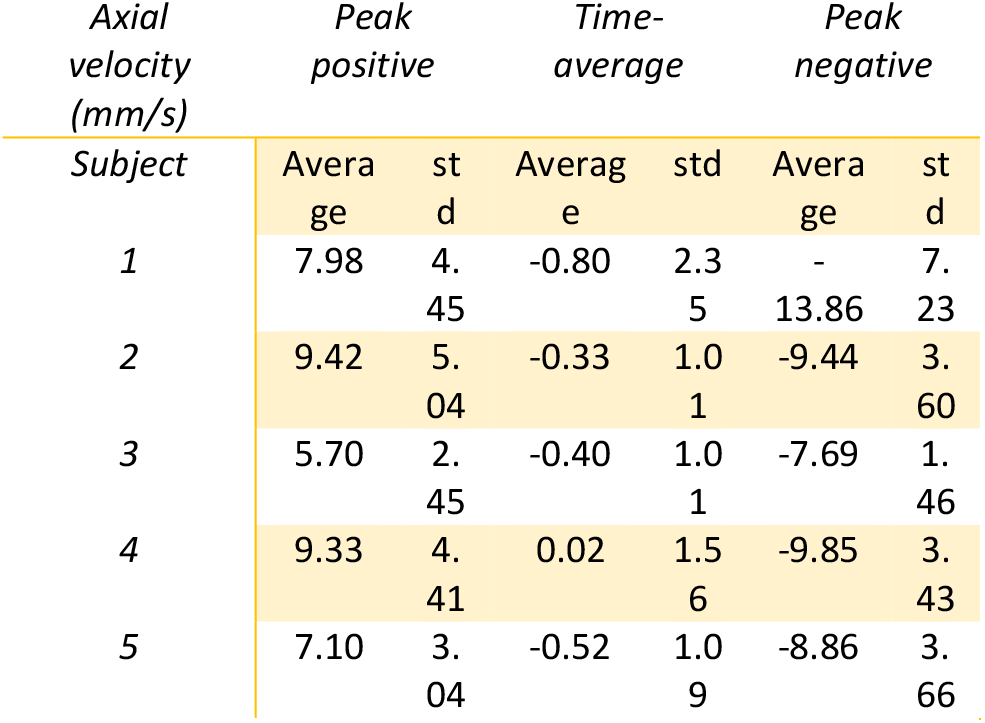
Axial velocity measurements of blood perfusion in the tibial cortex

**Table 4:**
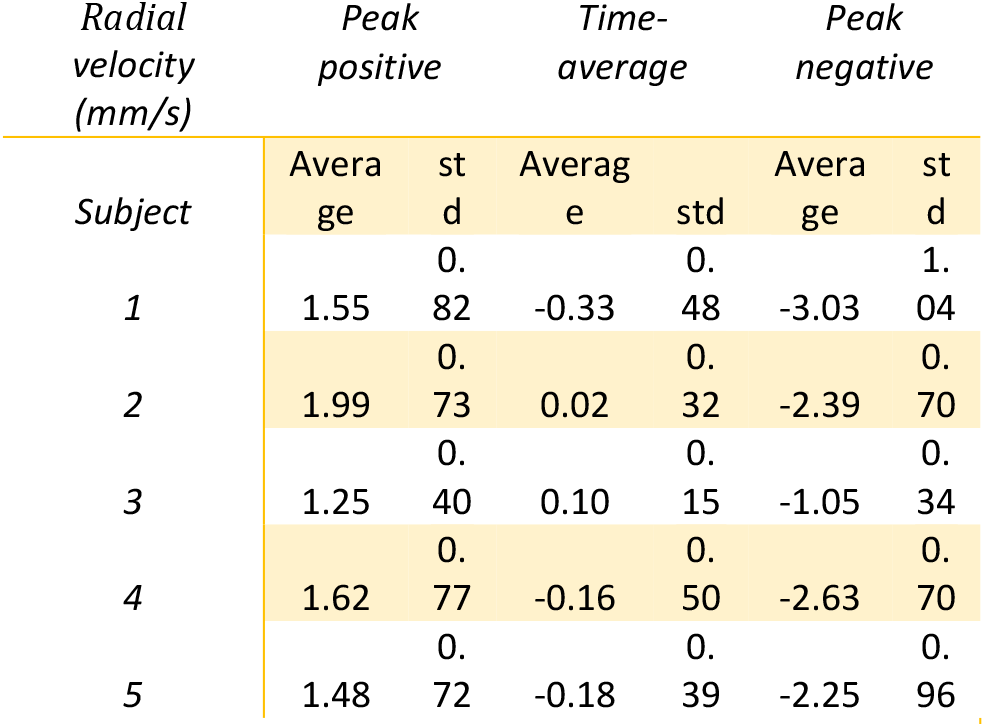
Radial velocity measurements of blood perfusion in the tibial cortex

The flow pulsatility frequency in cortical bone was measured very close to the heart rate observed in the superficial femoral artery, as expected (Table 5). However, no correlation was obtained between the flow peak velocity in the femoral artery and in cortical bone (not shown in the manuscript).

**Table 5:** Heart rate measured with the power Doppler signal in the superficial femoral artery and in tibial cortical bone

| Subject | Acquisition | Cardiac rate (beat/min) |  |
| --- | --- | --- | --- |
|  |  | Femoral artery | Cortical bone |
| 1 | 1 | 71 | 73 |
|  | 2 | 76 | 76 |
|  | 3 | 71 | 68 |
| 2 | 1 | 48 | 48 |
|  | 2 | 47 | 48 |
|  | 3 | 42 | 43 |
| 3 | 1 | 64 | 63 |
|  | 2 | 63 | 63 |
|  | 3 | 64 | 69 |
| 4 | 1 | 47 | 47 |
|  | 2 | 44 | 46 |
|  | 3 | 48 | 49 |
| 5 | 1 | 70 | 72 |
|  | 2 | 73 | 73 |
|  | 3 | 72 | 72 |

## Discussion

In this work, the first in vivo assessment of intraosseous blood circulation using ultrasound imaging was demonstrated. We proposed to assess blood perfusion in cortical bone simultaneously with femoral blood flow using plane-wave imaging and transverse oscillations in 5 healthy subjects. Pulsatile blood flow was observed in human cortical bone at the tibia with a pulsatility rate very close to the heart rate measured in the superficial femoral artery.

The axial velocity component was found 4 to 5 times larger than the radial velocity component suggesting a blood flow mainly in Haversian canals as expected from the intracortical vascular organization (9). In contrast, tibial cortical bone in mice possesses trans-cortical capillaries, mainly (14).

The axial and radial blood velocities measured in 5 healthy subjects are in the range expected for microvascular circulation, i.e., from 1 to 10 millimeters per second (40). Reproducible axial and radial velocity peaks were observed for every subject, indicating peak perfusion going from the heart to the foot and from the marrow to the cutaneous tissue (centrifugal flow). (41).

The centrifugal direction corroborates observations in animals and humans reported in the literature (41). In more than 90% of the cases, the human tibia has one nutrient artery, which penetrates the cortex through a nutrient foramen located below the tibial plateau, at the first proximal third of the tibial length (42). In the diaphyseal marrow of long bones, the nutrient artery divides into ascending and descending branches that supply the cortex with blood. However, the exact path taken by blood to circulate from the nutrient artery to the cortex of the diaphysis is poorly understood in humans (3,6). Our ultrasound examination was performed in the middle of the diaphysis. Interestingly we observed blood flow from the heart to the foot in all 5 volunteers, perhaps because the nutrient artery systematically enters the tibia at the first proximal third of the tibial length. The time-averaged blood perfusion was found to be less than 1 mm/s for both velocity components; therefore in agreement with perfusion rate measurements performed in animals with the microsphere technique, which suggested a mean blood velocity in the order of 1 mm/s for a purely unidirectional flow (16,43). Recently, blood velocity in the cortex of mice long bones was reported with intravital laser scanning confocal microscopy in the order of 1 mm/s (14). Thus our findings are in fair agreement with measurements reported with other technologies.

Even if the small number of subjects in this study does not allow us to infer general conclusions, it is interesting to note that the smallest peak velocity values were observed in the oldest subject (volunteer 3, age 65 y.o.). Based on the observation of cadaveric human long bones of different age, it was proposed that increasingly severe medullary ischemia with age, brought on by atherosclerosis of the marrow vessels, would cause blood supply of the cortex to evolve from a predominantly medullary blood supply to a predominantly periosteal blood supply (41). This evolution was thought to reduce the amount of circulating blood and its speed in the cortex The blood velocity was spatially averaged in a volume of tibial cortical bone of approximately 5mm x 15mm x 10mm. The image thickness (10mm) was determined by the out-of-plane width of the ultrasound beam generated by the phased-array probe used in this study. Clearly, the use of a matrix-array probe would improve the quantification of blood flow in the cortex because the information on moving blood could also be specified in the third spatial dimension. As a consequence, the hemodynamics parameters were likely underestimated because the true vascular organization deviates from our idealized description, assuming only two nearly perpendicular networks of parallel vessels. Moreover, the wide pores in cortical bone may host a small arteriole and a small venule. Our analysis likely integrates both arterial and venous flows in a resolution cell of approximately 1.5mm x 1.5mm in the ultrasound image, which leads to an underestimated measurement of cortical bone blood perfusion. Nonetheless, a pulsatile blood flow is expected in small arterioles only (44). The presented results demonstrate pulsatile blood flow in the tibial cortex, which suggests predominantly arterial blood circulation. This finding is in agreement with the fact that most canals contain a single vessel of capillary structure (7,15)

In this manuscript, an average wave-speed model for cortical bone was used for all subjects. Knowing that the compressional wave-speed in cortical bone can vary from one subject to another (45), we would expect a maximum error of 5% on the radial velocity component only. The femoral blood velocity assessment was essential to demonstrate that the pulsatility observed in the blood perfusion of cortical bone was trustful. However, due to the 4s recording, the simultaneous ultrasound imaging of the femoral artery and bone, and the hardware memory limitation, a frame rate of only 100 images per second has been used, which is the main limitation of the presented study. Indeed, although 100 images per second is enough to estimate a 10 mm/s blood flow, the limited frame rate and image number are not optimal for the SVD clutter filter. The quality and the spatial/temporal resolution of the measurements may be improved by increasing the number of tilted transmissions and the frame rate. The use of ultrasound contrast agents (46) may also significantly improve the estimated hemodynamic parameters’ sensitivity and specificity.

The presented approach was able to estimate very slow blood velocity (<5mm/s). Such blood velocity has already been estimated in the rat brain (40). The estimation of very slow blood velocities in a human bone was possible because bones are rigid, thus the only limitation is the motion of the bone relative to the probe, in particular out-of-plane motion. However, this limitation may be overcome by using a matrix-array probe that allows three-dimensional motion correction.

This manuscript presented results obtained at the diaphysis of the tibia; nonetheless, it could be easily applied to investigate the vascularization of other long bones such as the femur or radius. An interesting follow-up study could be the estimation of intraosseous blood circulation at a different location along a long bone to assess the heterogeneity of blood circulation in the cortex, which was demonstrated in animals with the microsphere technique (16).

The presented method might also be used to visualize and measure blood circulation in layers overlying and underlying the bone cortex such as the cutaneous tissue, muscle, marrow, and study the hemodynamic coupling between layers. The developed approach is expected to unlock the in vivo non-invasive assessment of intraosseous blood circulation in humans. Intraosseous ultrasonography could help to gain new knowledge on vascularization-related bone physio-pathological processes. It may help in the early diagnosis of bone diseases, in the monitoring of the effect of drugs or therapeutic treatments that have an action on intraosseous blood circulation, or in the monitoring of fracture healing.

## Appendix

## Ultrasonic imaging

We used a Vantage 256 ultrasound scanner (Verasonics, Kirkland, WA, USA). With 256 channels in emission and reception, we were able to connect two ultrasound transducers to the ultrasound scanner to image the femoral artery and the tibia simultaneously.

For imaging the femoral artery, we used a L7-4 linear array (ATL Phillips) composed of 128 piezoelectric elements spaced with a pitch of 298 μm. The probe has a central ultrasound frequency of 5 MHz. Four hundred ultrasound Doppler images at a frame rate of 100Hz were acquired. Each of them was obtained with the transmission of 30 identical titled plane waves (with a steering angle of 20°) at a pulse repetition rate of 5000 Hz.

For imaging the tibia, we used a P4-1 phased array (ATL Phillips) made of 96 piezoelectric elements spaced at a pitch of 295 μm. The probe has a central ultrasound frequency of 2.5 MHz. Four hundred high-resolution compound images at 100 Hz were acquired. Each high-resolution image was obtained from 15 planar insonifications tilted from −8° to +8° in the cutaneous tissue (Supplemental figure 1a).

## Refraction-corrected ultrasound image of the bone cortex

Intraosseous image reconstruction was based on an adaptation of the conventional delay-and-sum algorithm. The region of interest was regarded as a layered medium, and refraction occurring at the interface between layers was accounted for. Three layers are considered; cutaneous tissue, cortical bone, and marrow. Transmit and receive travel times were calculated with two-point ray tracing using Fermat’s principle with Brent’s method (32). The anisotropy of the compressional wave-speed in cortical bone was modeled with a model of weak transverse isotropy proposed by seismologists (47)

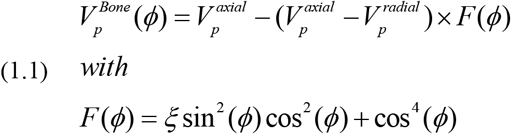

ϕ is the group angle, or acoustic ray angle (*ϕ*=0° means normal to the bone axis). 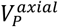 and 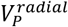 are the compressional wave-speeds in the direction of the bone axis and normal to it. ξ is an anisotropy shape parameter. In this work, we used the same wave-speed model for all subjects, with 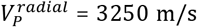, 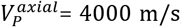, and ξ= 1.5. These values were estimated in two healthy volunteers in a previous study (32). The compressional wave-speed for the cutaneous tissue layer and the marrow layer was 1540 m/s and 1400 m/s, respectively.

The reconstruction algorithm sums the recorded echo signals 1) along the calculated round-trip travel times over the receive aperture of the probe array, and 2) over the steered plane wave transmissions (coherent compounding) (29).

## Measurement of blood flow in tibial cortical bone

The 2D vector field of blood velocity has been estimated in cortical bone using a method combining a Singular Value Decomposition (SVD) filter (34) and the transverse oscillation approach (38). The SVD filter was applied to the 400 high-contrast-resolution images. It removed the echo signals backscattered by the solid porous matrix of cortical bone. After the reception of ultrasonic echoes and image reconstruction, we obtain complex ultrasonic images s(x,z,t). After SVD processing, the spatiotemporal sequence corresponding to ultrasonic data can be rewritten as:

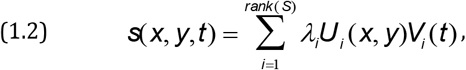

where S is the reshaped s matrix into the Casorati matrix with dimension (n_x_×n_y_, n_t_). Here U_i_ and V_i_ are the spatial and temporal singular vectors of the SVD decomposition. *λ_i_* are the ordered singular values based on|*λ_i_*|, *λ*_*i*+1_ < *λ_i_*. Then, the blood signal s_blood_ can be extracted using a threshold value n on the number of singular vectors as follows:

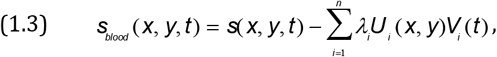

The SVD threshold value was chosen manually by looking at the corresponding power Doppler results for each subject (Table 1). Even if different SVD threshold values were used from one subject to another, the same SVD threshold has been used for the three acquisitions on the same subject. A significant improvement of the presented method could be using an automatic SVD threshold detection (48).

We calculated the power Doppler value *I* for each pixel as

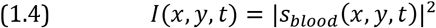

The transverse oscillation (TO) method (37–39) was used to calculate the axial and radial blood velocity. It aims to artificially introduce oscillations perpendicular to the ultrasound beam axis, i.e. in the direction of the long bone axis, to estimate the motion using a 2D phase-based approach. The s_blood_ image is convolved with the adapted filter to produce the transverse oscillations. In this study, we used the spatial transverse filter given in (4) made of the multiplication of a Gaussian window G(x) and a sinusoid

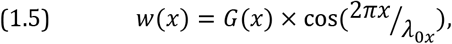

where x represents the position of the transducer element and λ_0x_ represents the desired TO wavelength, and we have

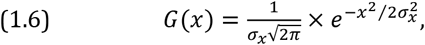

where σ_x_ is the width of the Gaussian, which is chosen to be of the same order as the ultrasound beam frequency spectrum’s width. The convolution is performed in the Fourier domain. The 2-D Fourier transform (FT) of the s_blood_ image is multiplied by a mask, Ω(λ_z_, λ_x_) where each line corresponds to the Fourier transform of ω(x):

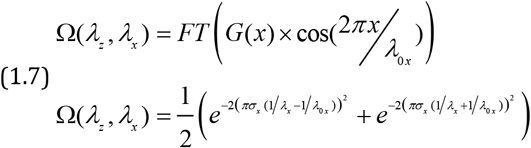

Once TO images have been obtained, it becomes possible to estimate the vector motion using the phase-based technique described in (49). This method aims to create complex image blocks based on a 2-D extension of the analytical signal using the Hann approach (50). Basically, an image block is generated by keeping only one quadrant of its 2-D Fourier spectrum. Two different analytical images are obtained by keeping two different quadrants of the 2-D Fourier spectra. The 2-D motion vector (*V*_axial_, *V*_lateral_) between two successive images is then deduced from the phases φ_11_, φ_12_, φ_21_, and φ_22_ according to

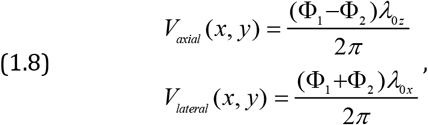

with

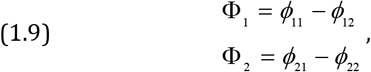

where λ_0z_ is the ultrasound beam wavelength of s_blood_ images corresponding to the wavelength of the signal received by the probe, *ϕ*_11_ and *ϕ*_12_ are the phases of two analytical images from the first image, and *ϕ*_21_ and *ϕ*_22_ are the phases of two analytical images from the next consecutive image. Note that the analytical phase images were smoothed in space and time using a moving average filter of 5 time samples (50 ms) and 5 per 5 spatial samples (1.5 mm × 1.5 mm). For further detail on this motion estimation technique, the reader can refer to (49). In this study, the same λ_0z_ and λ_0x_ values have been used for every subject and acquisition. λ_0z_ was taken equal to 1.3mm by assuming a speed of sound of 3250 m/s in the cortical bone (32,45) and with a central transducer frequency of 2.5MHz. λ_0x_ was chosen empirically and set to 3.5mm. Finally, for each image pixel, the radial and axial velocity time curves were filtered with an in-house Fourier series selection method by retaining only the 5 frequency components with the highest amplitude (an example is shown in Supplemental figure 2). The TO implementation and velocity estimation codes and filtering method are provided as Supplementary Information.

## Measurement of blood flow in the superficial femoral artery

Each ensemble of 30 high frame rate images has been filtered using the same SVD approach, but with a threshold value of 4. Then, the conventional Doppler technique was applied to each ensemble. The blood velocity in the direction of the artery axis was calculated using:

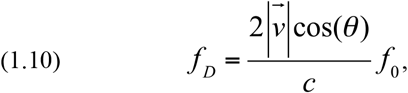

where *f*_D_ and *f*_D_ are the “Doppler frequency” and the ultrasound frequency, respectively, 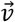 is the blood velocity, and cis the speed of sound. The beam-to-flow angle *θ* was manually determined in B-mode images.

## Heart rate estimation

Because the power Doppler signal is more robust to noise, we used this hemodynamic parameter to estimate the heart rate. The cardiac rate was measured automatically by finding the local maxima of the temporal power Doppler signal.

The cardiac rate varied from one acquisition to another (Table 5). Consequently, 3 average cardiac rates corresponding to the 3 acquisitions have been measured for each subject. Nonetheless, a good correlation was found between the femoral artery and the cortical bone pulsatility rate for each acquisition and subject.

## Code availability

The code for image reconstruction of the bone cortex is available as supplementary material of reference 33. The code for applying the transverse oscillation method is available online at https://www.creatis.insa-lyon.fr/ius-special-issue-2014/ (see also reference 39). Custom scripts in MATLAB for filtering of the velocity signals (Fourier series selection) are available in the **Supplementary Information**.

## Data availability

The authors declare that all data supporting this study’s findings are available within the paper and its Supplementary Information. Raw acquired ultrasound data can be made available upon reasonable request, with permission of Erasmus MC, The Netherlands, and Sorbonne Université, France.

## Acknowledgment

This work has been funded by a French ANR program (project FUIBCA ANR-17-CE19-0008-01) and by SATT LUTECH (project BONES).

## Supplementary material

**Supplemental Figure 1:**
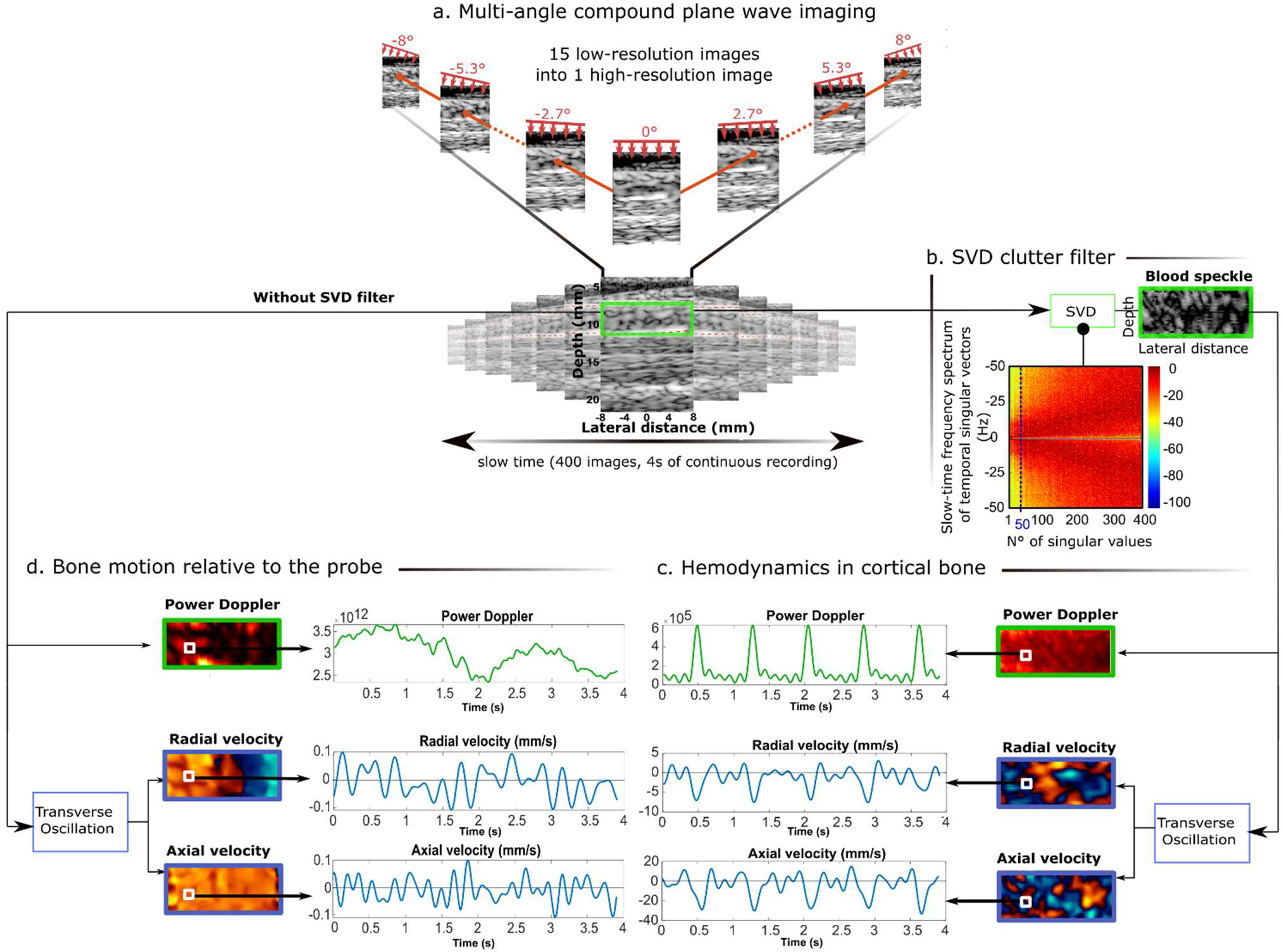
Workflow for the estimation of intraosseous blood perfusion. a) Ultrasound imaging of bone was performed by acquiring 400 compound images at a frame rate of 100 Hz, each frame obtained from 15 planar illuminations tilted from −8° to +8° in the cutaneous tissue. b) The blood signal was extracted from the bone signal by applying a clutter filter based on singular value decomposition (SVD). c) The power Doppler was calculated from the resulting sequences. Moreover, the axial (long bone axis) and radial (trans-cortex) 1components of the blood velocity were estimated using the transverse oscillation approach. d) The same procedure applied to the raw reconstructed images (without SVD filter) shows bone motion relative to the hand-held probe. The hemodynamic parameters are shown for a single pixel indicated in the images by a white square.

**Supplemental Figure 2:**
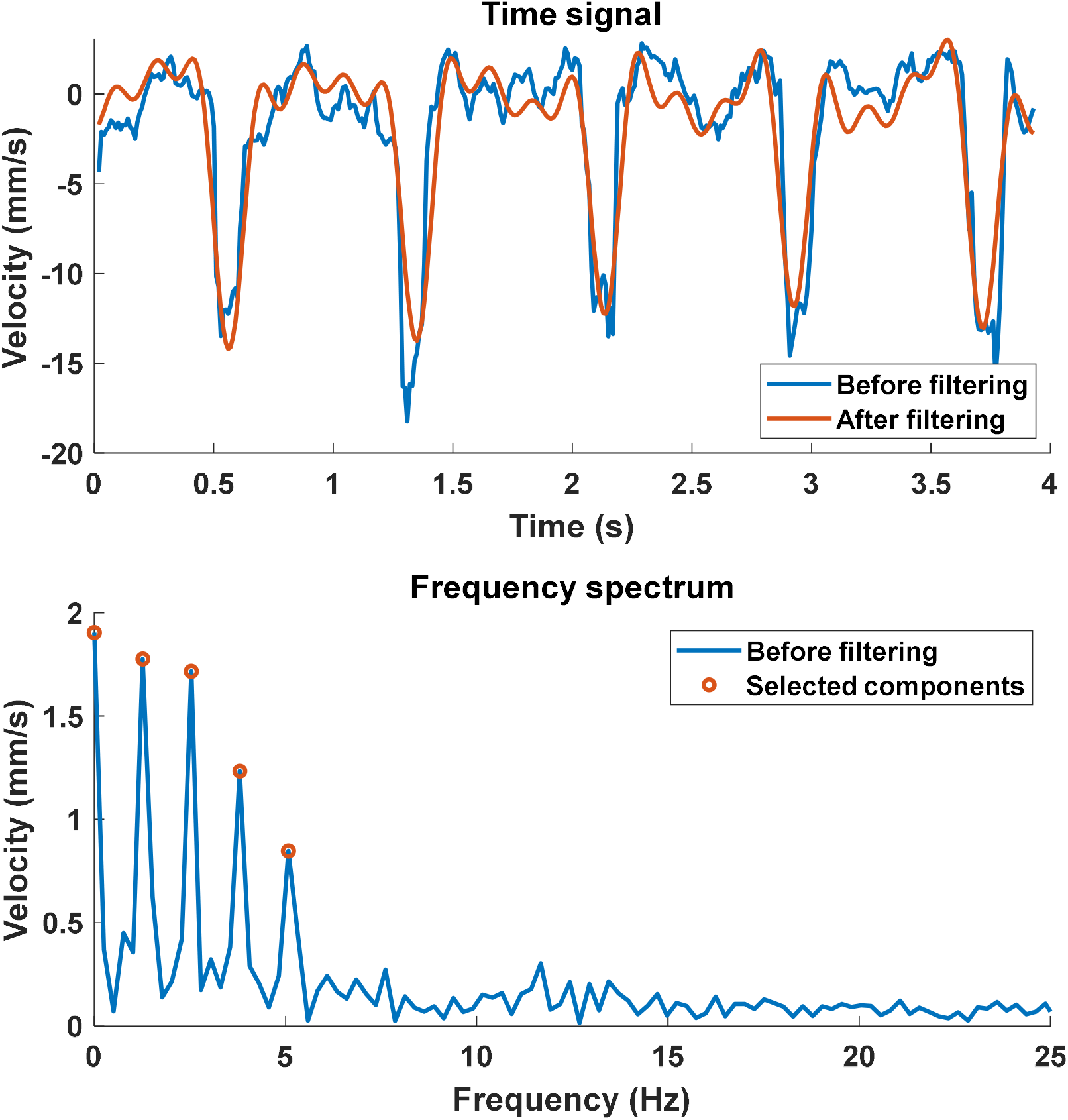
Fourier series selection filtering method. In the top panel, the blue and red curves are obtained before and after filtering, respectively. The 5 selected frequencies are indicated with red circles in the bottom panel.

## References

1. Ramasamy SK., Kusumbe AP., Schiller M., et al. Blood flow controls bone vascular function and osteogenesis. Nat Commun 2016;7(1):13601. Doi: 10.1038/ncomms13601.

2. Cowin SC., Cardoso L. Blood and interstitial flow in the hierarchical pore space architecture of bone tissue. Journal of Biomechanics 2015;48(5):842–54. Doi: 10.1016/j.jbiomech.2014.12.013.

3. Lafage-Proust M-H., Roche B., Langer M., et al. Assessment of bone vascularization and its role in bone remodeling. BoneKEy Reports 2015;4. Doi: 10.1038/bonekey.2015.29.

4. Lafage-Proust M-H., Prisby R., Roche B., Vico L. Bone vascularization and remodeling. Joint Bone Spine 2010;77(6):521–4. Doi: 10.1016/j.jbspin.2010.09.009.

5. Tomlinson RE., Silva MJ. Skeletal Blood Flow in Bone Repair and Maintenance. Bone Res 2013;1(4):311–22. Doi: 10.4248/BR201304002.

6. Marenzana M., Arnett TR. The Key Role of the Blood Supply to Bone. Bone Res 2013;1(3):203–15. Doi: 10.4248/BR201303001.

7. Laroche M. Intraosseous circulation from physiology to disease. Joint Bone Spine 2002;69(3):262–9. Doi: 10.1016/S1297-319X(02)00391-3.

8. Alagiakrishnan K., Juby A., Hanley D., Tymchak W., Sclater A. Role of Vascular Factors in Osteoporosis. The Journals of Gerontology Series A: Biological Sciences and Medical Sciences 2003;58(4):M362–6. Doi: 10.1093/gerona/58.4.M362.

9. Maggiano IS., Maggiano CM., Clement JG., Thomas CDL., Carter Y., Cooper DML. Three-dimensional reconstruction of Haversian systems in human cortical bone using synchrotron radiation-based micro-CT: morphology and quantification of branching and transverse connections across age. J Anat 2016;228(5):719–32. Doi: 10.1111/joa.12430.

10. Hennig C., Thomas CDL., Clement JG., Cooper DML. Does 3D orientation account for variation in osteon morphology assessed by 2D histology? J Anat 2015;227(4):497–505. Doi: 10.1111/joa.12357.

11. Cooper DML., Thomas CDL., Clement JG., Turinsky AL., Sensen CW., Hallgrímsson B. Age-dependent change in the 3D structure of cortical porosity at the human femoral midshaft. Bone 2007;40(4):957–65. Doi: 10.1016/j.bone.2006.11.011.

12. Thomas CDL., Feik SA., Clement JG. Increase in pore area, and not pore density, is the main determinant in the development of porosity in human cortical bone. J Anat 2006;209(2):219–30. Doi: 10.1111/j.1469-7580.2006.00589.x.

13. Stein MS., Feik SA., Thomas CD., Clement JG., Wark JD. An automated analysis of intracortical porosity in human femoral bone across age. J Bone Miner Res 1999;14(4):624–32. Doi: 10.1359/jbmr.1999.14.4.624.

14. Grüneboom A., Hawwari I., Weidner D., et al. A network of trans-cortical capillaries as mainstay for blood circulation in long bones. Nat Metab 2019;1(2):236–50. Doi: 10.1038/s42255-018-0016-5.

15. Brookes M., Revell WJ. Blood Supply of Bone. London: Springer London; 1998.

16. McCarthy I. The Physiology of Bone Blood Flow: A Review. J Bone Joint Surg Am 2006;88(suppl_2):4. Doi: 10.2106/JBJS.F.00890.

17. Serrat MA. Measuring bone blood supply in mice using fluorescent microspheres. Nat Protoc 2009;4(12):1749–58. Doi: 10.1038/nprot.2009.190.

18. Estrada H., Rebling J., Sievert W., et al. Intravital optoacoustic and ultrasound bio-microscopy reveal radiation-inhibited skull angiogenesis. Bone 2020;133:115251. Doi: 10.1016/j.bone.2020.115251.

19. Dyke JP., Aaron RK. Noninvasive methods of measuring bone blood perfusion: Noninvasive methods of measuring bone blood perfusion. Annals of the New York Academy of Sciences 2010;1192(1):95–102. Doi: 10.1111/j.1749-6632.2009.05376.x.

20. Griffith JF., Yeung DK., Tsang PH., et al. Compromised Bone Marrow Perfusion in Osteoporosis. J Bone Miner Res 2008;23(7):1068–75. Doi: 10.1359/jbmr.080233.

21. Baur-Melnyk A. Magnetic Resonance Imaging of the Bone Marrow. Berlin, Heidelberg: Springer Berlin Heidelberg; 2013.

22. Wan L., Wu M., Sheth V., et al. Evaluation of cortical bone perfusion using dynamic contrast enhanced ultrashort echo time imaging: a feasibility study. Quant Imaging Med Surg 2019;9(8):1383–93. Doi: 10.21037/qims.2019.08.05.

23. Ashcroft GP., Evans NT., Roeda D., et al. Measurement of blood flow in tibial fracture patients using positron emission tomography. The Journal of Bone and Joint Surgery British Volume 1992;74(5):673–7.

24. Binzoni T., Spinelli L. Near-infrared photons: a non-invasive probe for studying bone blood flow regulation in humans. J Physiol Anthropol 2015;34(1):28. Doi: 10.1186/s40101-015-0066-2.

25. Jensen JA., Nikolov S., Yu ACH., Garcia D. Ultrasound Vector Flow Imaging: I: Sequential Systems. IEEE Trans Ultrason, Ferroelect, Freq Contr 2016:1–1. Doi: 10.1109/TUFFC.2016.2600763.

26. Jensen JA., Nikolov SI., Yu ACH., Garcia D. Ultrasound Vector Flow Imaging—Part II: Parallel Systems. IEEE Trans Ultrason, Ferroelect, Freq Contr 2016;63(11):1722–32. Doi: 10.1109/TUFFC.2016.2598180.

27. Sun M-H., Leung K-S., Zheng Y-P., et al. Three-dimensional high frequency power Doppler ultrasonography for the assessment of microvasculature during fracture healing in a rat model: 3D HIGH FREQUENCY POWER DOPPLER ULTRASONOGRAPHY. J Orthop Res 2012;30(1):137–43. Doi: 10.1002/jor.21490.

28. Doria AS., Guarniero R., Cunha FG., et al. Contrast-enhanced power Doppler sonography: assessment of revascularization flow in Legg-Calvé-Perthes’ disease. Ultrasound in Medicine & Biology 2002;28(2):171–82. Doi: 10.1016/S0301-5629(01)00500-2.

29. Montaldo G., Tanter M., Bercoff J., Benech N., Fink M. Coherent plane-wave compounding for very high frame rate ultrasonography and transient elastography. IEEE Trans Ultrason Ferroelectr Freq Control 2009;56(3):489–506. Doi: 10.1109/TUFFC.2009.1067.

30. Udesen J., Gran F., Hansen KL., Jensen JA., Thomsen C., Nielsen MB. High frame-rate blood vector velocity imaging using plane waves: simulations and preliminary experiments. IEEE Trans Ultrason Ferroelectr Freq Control 2008;55(8):1729–43. Doi: 10.1109/TUFFC.2008.858.

31. Bercoff J., Montaldo G., Loupas T., et al. Ultrafast compound doppler imaging: providing full blood flow characterization. IEEE Trans Ultrason, Ferroelect, Freq Contr 2011;58(1):134–47. Doi: 10.1109/TUFFC.2011.1780.

32. Renaud G., Kruizinga P., Cassereau D., Laugier P. In vivo ultrasound imaging of the bone cortex. Phys Med Biol 2018;63(12):125010. Doi: 10.1088/1361-6560/aac784.

33. Renaud G., Clouzet P., Cassereau D., Talmant M. Measuring anisotropy of elastic wave velocity with ultrasound imaging and an autofocus method - application to cortical bone. Phys Med Biol 2020. Doi: 10.1088/1361-6560/abb92c.

34. Demene C., Deffieux T., Pernot M., et al. Spatiotemporal Clutter Filtering of Ultrafast Ultrasound Data Highly Increases Doppler and fUltrasound Sensitivity. IEEE Trans Med Imaging 2015;34(11):2271–85. Doi: 10.1109/TMI.2015.2428634.

35. Rubin JM., Bude RO., Carson PL., Bree RL., Adler RS. Power Doppler US: a potentially useful alternative to mean frequency-based color Doppler US. Radiology 1994;190(3):853–6. Doi: 10.1148/radiology.190.3.8115639.

36. Yiu BYS., Yu ACH. Least-Squares Multi-Angle Doppler Estimators for Plane-Wave Vector Flow Imaging. IEEE Trans Ultrason, Ferroelect, Freq Contr 2016;63(11):1733–44. Doi: 10.1109/TUFFC.2016.2582514.

37. Jensen JA., Munk P. A new method for estimation of velocity vectors. IEEE Trans Ultrason Ferroelectr Freq Control 1998;45(3):837–51. Doi: 10.1109/58.677749.

38. Salles S., Chee AJY., Garcia D., Yu ACH., Vray D., Liebgott H. 2-D arterial wall motion imaging using ultrafast ultrasound and transverse oscillations. IEEE Trans Ultrason Ferroelectr Freq Control 2015;62(6):1047–58. Doi: 10.1109/TUFFC.2014.006910.

39. Salles S., Liebgott H., Garcia D., Vray D. Full 3-D transverse oscillations: a method for tissue motion estimation. IEEE Transactions on Ultrasonics, Ferroelectrics, and Frequency Control 2015;62(8):1473–85. Doi: 10.1109/TUFFC.2015.007050.

40. Macé E., Montaldo G., Cohen I., Baulac M., Fink M., Tanter M. Functional ultrasound imaging of the brain. Nat Methods 2011;8(8):662–4. Doi: 10.1038/nmeth.1641.

41. Bridgeman G., Brookes M. Blood supply to the human femoral diaphysis in youth and senescence. J Anat 1996;188 (Pt 3):611–21.

42. Kirschner MH., Menck J., Hennerbichler A., Gaber O., Hofmann GO. Importance of arterial blood supply to the femur and tibia for transplantation of vascularized femoral diaphyses and knee joints. World J Surg 1998;22(8):845–51; discussion 852. Doi: 10.1007/s002689900480.

43. Anetzberger H., Birkenmaier C. THE MICROSPHERE METHOD FOR INVESTIGATING BONE BLOOD FLOW. Skeletal Circulation in Clinical Practice. WORLD SCIENTIFIC; 2016. p. 53–84.

44. Santisakultarm TP., Cornelius NR., Nishimura N., et al. In vivo two-photon excited fluorescence microscopy reveals cardiac- and respiration-dependent pulsatile blood flow in cortical blood vessels in mice. Am J Physiol Heart Circ Physiol 2012;302(7):H1367–1377. Doi: 10.1152/ajpheart.00417.2011.

45. Eneh CTM., Malo MKH., Karjalainen JP., Liukkonen J., Töyräs J., Jurvelin JS. Effect of porosity, tissue density, and mechanical properties on radial sound speed in human cortical bone: Radial sound speed in human cortical bone. Med Phys 2016;43(5):2030–9. Doi: 10.1118/1.4942808.

46. Hingot V., Errico C., Heiles B., Rahal L., Tanter M., Couture O. Microvascular flow dictates the compromise between spatial resolution and acquisition time in Ultrasound Localization Microscopy. Sci Rep 2019;9(1):2456. Doi: 10.1038/s41598-018-38349-x.

47. Thomsen L. Weak elastic anisotropy. GEOPHYSICS 1986;51(10):1954–66. Doi: 10.1190/1.1442051.

48. Baranger J., Arnal B., Perren F., Baud O., Tanter M., Demene C. Adaptive Spatiotemporal SVD Clutter Filtering for Ultrafast Doppler Imaging Using Similarity of Spatial Singular Vectors. IEEE Trans Med Imaging 2018;37(7):1574–86. Doi: 10.1109/TMI.2018.2789499.

49. Liebgott H., Basarab A., Gueth P., Friboulet D., Delachartre P. Transverse oscillations for tissue motion estimation. Ultrasonics 2010;50(6):548–55. Doi: 10.1016/j.ultras.2009.11.001.

50. Basarab A., Gueth P., Liebgott H., Delachartre P. Phase-based block matching applied to motion estimation with unconventional beamforming strategies. IEEE Trans Ultrason, Ferroelect, Freq Contr 2009;56(5):945–57. Doi: 10.1109/TUFFC.2009.1127.

